# Hemin-induced platelet activation is regulated via ACKR3 chemokine surface receptor - implications for passivation of vulnerable atherosclerotic plaque

**DOI:** 10.1101/2024.05.13.593847

**Authors:** Zoi Laspa, Valerie Dicenta-Baunach, David Schaale, Manuel Sigle, Ravi Hochuli, Tatsiana Castor, Alp Bayrak, Tobias Harm, Karin Anne Lydia Mueller, Thanigaimalai Pillaiyar, Stefan Laufer, Anne-Katrin Rohlfing, Meinrad Paul Gawaz

## Abstract

In vulnerable atherosclerotic plaques intraplaque hemorrhages (IPH) result in hemolysis of red blood cells and release of hemoglobin and free hemin. Hemin activates platelets and leads to thrombosis. Agonism of the inhibitory platelet receptor ACKR3 inhibits hemin-dependent platelet activation and thrombus formation.

To characterize the effect of hemin and ACKR3 agonism on isolated human platelets, multi-color flow cytometry and classical experimental setup such as light transmission aggregometry and a flow chamber assay have been used.

Hemin induces platelet aggregation and *ex vivo* platelet-dependent thrombus formation on immobilized collagen under low shear rate 500 s^-1^ indicating that free hemin is a strong activator for platelet-dependent thrombosis. Recently, we described that ACKR3 is a prominent inhibitory receptor of platelet activation. Specific ACKR3 agonists but not conventional antiplatelet compounds such as COX-1 inhibitor (indomethacin), ADP-receptor blocker (cangrelor), or PAR1 inhibitor (ML161) inhibit both hemin-dependent aggregation and thrombus formation. To further characterize the effect of hemin on platelet subpopulations we established a multi-color flow cytometry assay. We found that hemin induces procoagulant (CD42b^pos^/ PAC-1^neg^/ AnnexinV^pos^), aggregatory (CD42b^pos^ / PAC-1^pos^ / AnnexinV^neg^) and inflammatory (CD42b^pos^/ CXCR4^pos^/ACKR3^pos^/ AnnexinV^pos^) platelet subpopulations. Treatment with ACKR3 agonists significantly decrease the formation of procoagulant and ACKR3^pos^ platelets in response to hemin.

We conclude that hemin is a strong activator for the formation of procoagulant platelets and thrombus formation which is dependent on the function of ACKR3. Activation of ACKR3 through specific agonists may offer a therapeutic strategy to control vulnerability of atherosclerotic plaques in areas of IPH.

**Graphical abstract:** Intraplaque hemorrhages (IPH) results in hemolysis and liberation of iron containing heme and its oxidized metabolite hemin. Hemin activates platelets and has strong pro-thrombotic activity. Agonism of the atypical chemokine receptor 3 (ACKR3) inhibits hemin-induced platelet activation.

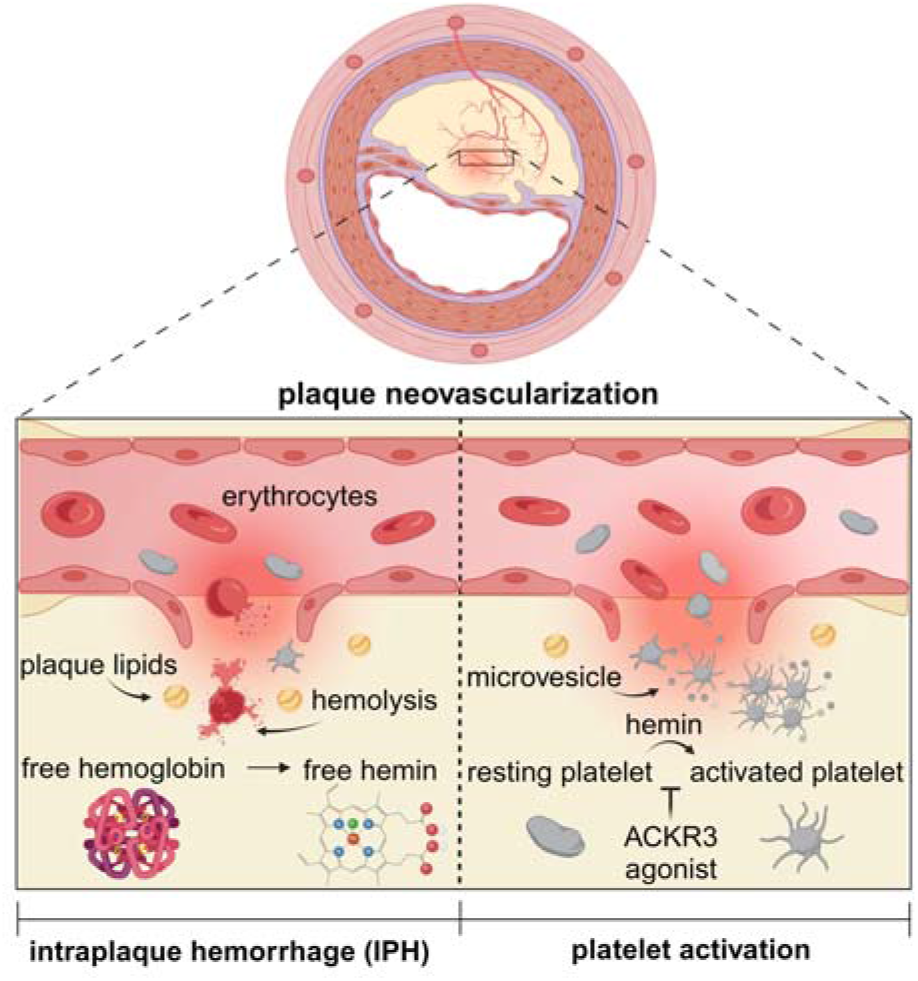

## Introduction

Vulnerability of atherosclerotic plaques is associated with an increased risk for myocardial infarction and ischemic stroke [1-7]. Instability of a vulnerable plaque is characterized by enhanced lipid accumulation (lipid core), inflammation and thrombus formation [5-7]. Intraplaque hemorrhages result from rupture of fragile neovessels within the atherosclerotic plaque, leading to bleeding into the plaque itself. Over time, this hemorrhage can trigger inflammation and thrombosis, exacerbating plaque instability and potentially leading to plaque rupture, hence intraplaque hemorrhages (IPH) have been recognized as a major factor for atheroprogression and plaque vulnerability and is found in up to 40% of high-risk plaques and has been identified as trigger for thrombo-ischemic clinical events [8-13]. Upon IPH red blood cells (RBCs) accumulate extracellularly in the diseased vascular tissue [14]. Interaction of RBCs and plaque lipids results in erythrocyte lysis and release of free hemoglobin (Hb) [15]. Degradation of extracellular Hb outside the protective environment of RBCs results in liberation of free iron-containing heme and its oxidized metabolite hemin [15, 16]. Free heme elicits prothrombotic and proinflammatory activities [17].

Recently, hemin has been shown to induce platelet activation via interaction of C-type lectin-like receptor 2 (CLEC-2) and glycoprotein VI (GPVI) [18, 19]. Hemin induces platelet plasma disintegration and shedding of membrane receptors including GPVI and P-selectin (CD62P) [20, 21]. Hemin-dependent platelet activation and platelet-dependent thrombus formation is regulated via the subtilisin-like proprotein convertase furin and the second messenger cGMP [21, 22]. Conventional antiplatelet drugs are insufficient to mitigate hemin-dependent platelet activation [21]. Only recently, we have defined platelet ACKR3 (formally known as CXCR7) as inhibitory receptor on platelets [23-25]. ACKR3 agonism inhibits platelet activation both *in vitro* and an *in vivo* mouse model of experimental myocardial infarction [24].

The purpose of the present study was to further characterize the impact of hemin on platelet functions and the multitudinous pattern of platelet phenotypes. We found that hemin induces substantial formation of procoagulant platelets and microvesicles [21]. The hemin-dependent generation of procoagulant platelets can be attenuated in the presence of specific ACKR3 agonists but not with conventional platelet inhibitors. The findings imply that ACKR3 agonists may provide a therapeutic tool to control vulnerability of atherosclerotic plaques in areas of IPH.

## Methods

### Chemicals and antibodies

Monoclonal mouse anti-CXCR7 BV421 conjugated antibody, monoclonal mouse anti-human CD184 BV650 conjugated antibody, monoclonal mouse anti-human CD61 BV605 conjugated antibody and monoclonal anti-human PAC-1 FITC antibody were obtained from BD Bioscience (Franklin Lakes, NJ, USA). Monoclonal mouse anti-CD63 PE/Cyanine5 antibody was obtained from Abcam (Cambridge, UK). Monoclonal mouse anti-human CD42b PerCP/Cyanine5.5 conjugated antibody, Alexa Fluor® 700 anti-human CD41 antibody and Zombie NIR^®^ fixable dye were obtained from BioLegend (San Diego, CA, USA). Anti-human CD62P conjugated PE antibody was obtained from Beckman Coulter (Brea, California, USA). Donkey anti-rabbit IR-Dye 680RD and donkey anti-rabbit IR-Dye 800CW were obtained from Li-COR (Lincoln, USA). Phalloidin Alexa Fluor® 488, Fluo-4 and Annexin V PE-Cyanine7 were obtained from Invitrogen (Carlsbad, California, USA). Hemin, indometacin (COX-1 inhibitor), 3,3’
s-dihexyloxacarbocyanine iodine (DiOC_6_) and fibrinogen from human plasma were obtained from Sigma Aldrich (St. Louis, Missouri, USA). ML161 (PAR1 inhibitor) was obtained from Tocris (Bristol, UK). Cangrelor (KENGREXAL® 50 mg) was obtained from Chiesi Farmaceutici (Parma, Italy). CRP-XL (collagen related peptide) was obtained from CambCol Laboratories (Ely, UK).

VUF11207 (ACKR3 agonist) was obtained from Merck (Darmstadt, Germany). BY-061 and control compound C46 were described previously [24, 31].

### Isolation of human platelets

Blood was collected from healthy donors. Participants gave written informed consent for blood collection and the procedure was approved by Ethics Committee at the Medical Faculty of the Eberhard Karls University and at the University Hospital of Tuebingen (ethics vote 238/2018B02). Platelet isolation was performed as described previously [24, 39]. Blood was assembled in syringes containing acid-citrate-dextrose (ACD) anticoagulant (1:5) and centrifuged for 20 min at 209 xg without brake. The resulting platelet rich plasma (PRP) supernatant was added to Tyrode’
ss buffer pH 6.5 (137 mM NaCl, 2.8 mM KCl, 12 mM NaHCO_3_, 5 mM glucose, 10 mM HEPES). Next, the blood was centrifuged for 10 min at 836 xg with brake. The supernatant was discarded, the platelet pellet was resuspended in Tyrode’s buffer pH 7.4 and the platelet count was measured using a conventional cell counter (Sysmex Coorporation, Kobe, Japan).

### Multicolor Platelet Flow cytometry

Standard flow cytometry experiments were performed as described before [20]. Isolated human platelets (1×10^6^ per sample) were pre-incubated with ACKR3 agonists (100 µM VUF11207, 100 µM BY-061, 100 µM C46) for 15 min at room temperature (RT). Hemin and CRP-XL at indicated concentrations were added and samples were incubated with fluorochrome conjugated antibodies (anti-CD62P PE, anti-PAC-1 FITC, anti-CD63 PE/Cy5) for 30 min at room temperature. Afterwards the cells were fixed with 0.5% formaldehyde and analyzed with a flow cytometer (FACS Calibur flow cytometer, BD Bioscience, Franklin Lakes, NJ, USA). FlowJo (Version 10.8.0 FlowJo CCL, Ashton, OR, USA) was used to analyze the raw data.

For multicolor flow cytometry isolated human platelets were treated as described above. Samples with 1×10^6^ isolated human platelets were prepared in binding buffer (Invitrogen, Carlsbad, CA, USA). Hemin and CPR-XL were incubated at the indicated concentrations for 1h at room temperature.

Fluorescent-labeled antibodies (anti-CD42b PerCP/Cyanine5.5, anti-CD61 BV605, anti-CD41 Alexa Fluor^®^ 700, anti-CD63 PE/Cy5, anti-CD62P PE, anti-PAC-1 FITC, anti-CXCR4 BV650, anti-CXCR7 BV421), Annexin V PE-Cyanine7 and the fluorescent dye Zombie NIR were incubated for 30 min at room temperature. After that cells were diluted in 300 µl binding buffer and samples were analyzed immediately by flow cytometry (BD LSRFortessa™, BD Bioscience, San Jose, CA, USA). OMIQ flow cytometry software (Dotmatics, Boston, MA, USA) was used to analyze the raw data.

To quantify platelet-derived microvesicles isolated human platelets were treated as described under the standard flow cytometry protocol. Hemin and CRP-XL were incubated at indicated concentrations for 1h at room temperature. Then, samples were fixed with 0.5% formaldehyde and FSC/SSC data collected for 45 s using a FACSLyric™ (BD Bioscience, San Jose, CA, USA). Additionally, a suspension with 1 µm beads was measured to identify the microvesicular range of 1 µm within the FSC parameter.

Image stream analysis of platelets were performed using an anti-CXCR7 BV421 and an anti-CD42b PE antibody [40]. Washed human platelets (1×10^6^ per sample) were incubated with hemin (6.25 or 25 µM) or CRP-XL (1 µg/ml) for 30 min at room temperature. Subsequently, cells were diluted in 300 µl binding buffer and centrifuged at 400 xg for 5 min at room temperature. The supernatant was removed and the platelet pellet was resuspended in binding buffer. The cell suspension was analyzed immediately with an ImageStream^X^ Mark II Imaging Flow Cytometer (amnis, Seattle, WA, USA) and a 60x magnification. CD42b^pos^ cells were gated and the MFI of CXCR7 was used for representative images for every condition. Data were analyzed with IDEAS^®^ Image analysis software (Version 6.3).

### Platelet light transmission aggregometry

Light transmission aggregometry (CHRONO-LOG Aggregometer 490-X, Chrono-log Corporation, Havertown, PA, USA) was performed with isolated human platelets (4 x 10^7^ per sample) supplemented with 2 mM CaCl_2_ and 100 µg/ml fibrinogen [24]. The platelets were pre-incubated with ACKR3 agonists (100 µM VUF11207, 100 µM BY-061, 100 µM C46) or 10 µM indomethacin/ cangrelor/ PAR1 inhibitor ML161 for 15 min at 37°C. Aggregation was induced by hemin (6.25 µM) and maximal aggregation and the area under the curve was measured for 5:30 min.

### Platelet spreading

Coverslips were coated with fibrinogen (100 µg/ml) and incubated at 4°C overnight. The following day, the coverslips were washed with Dulbecco’s phosphate buffered solution (PBS) and isolated platelets were adjusted to 20,000/µl. Platelets were pre-incubated with ACKR3 agonist (100 µM VUF11207) for 15 min at room temperature. The samples were supplemented with 1 mM CaCl_2_ and incubated with hemin on fibrinogen coated coverslips for 30 min. After that, the platelets were fixed for 15 min with 2% formaldehyde and permeabilized with 0.1% Triton-X-100 in PBS for 5 min. Afterward, Phalloidin-Alexa488 (1:200) was added and incubated for 1 h at room temperature. The coverslips were mounted onto slides and five images at randomly selected areas were taken with a Nikon Eclipse Ti2 microscope (Nikon Instruments Europe BV, Amsterdam, The Netherlands). The images were analyzed with the NIS-Elements AR software (Version 5.21.00, Nikon Instruments Europe BV, Amsterdam, The Netherlands).

### Calcium release measurement

The intracellular Ca^2+^ - release induced by hemin was measured by Fluo4 fluorescence intensity with a plate reader (GloMax®-Multi Detection System, Promega GmbH, Walldorf, Germany). Isolated platelets (6 x 10^7^ per sample) were pre-incubated with 5 µM Fluo4 for 30 min at room temperature. After that, the samples were incubated with an ACRK3 agonist (100 µM VUF11207) or Tyrode’
ss buffer as vehicle control for 15 min at room temperature. Subsequently, platelets were activated with hemin (6.25 µM) and the fluorescence was measured manually at 0 s and 30 s. Afterward, the fluorescence was automatically measured every 60 s over a time period of 5 min.

### Flow chamber assay

For *in vitro* thrombus formation coverslips were coated with 100 µg/ml collagen (Collagen Reagens HORM® Suspension, Takeda Pharmaceutical Company, Tokyo, Japan) and blocked with 1% bovine serum albumin (BSA) in PBS containing Ca^2+^ at room temperature. Washed platelets (4 x 10^8^ per sample) in PBS with Ca^2+^ (5:1) supplemented with 1 mM CaCl_2_ were incubated with 1 µM 3,3’-Dihexyloxacarbocyanine Iodide (DiOC_6_) for 10 min and ACKR3 agonists (100 µM VUF11207, 100 µM C46) for 15 min at room temperature in the dark. Then, samples were stimulated with 6.25 µM hemin and the thrombus formation at a low shear rate 500 s^-1^ was visualized in a flow chamber setup (Maastricht Instruments B.V., Maastricht, The Netherlands) using a Nikon Eclipse Ti2-A microscope. After 10 min five images were taken randomly and the thrombus formation was measured as area fraction covered by the platelets and analyzed with NIS-Elements AR software (Version 5.21.00, Nikon Instruments Europe BV, Amsterdam, The Netherlands).

### Immunoblot analysis

Isolated human platelets (2.5×10^8^ per sample) were activated with hemin (6.25 or 25 µM) for 5 min at room temperature. Cells were lysed in RIPA lysis buffer (50 mM TRIS/HCl, 150 mM NaCl, 0.1% SDS, 1% Triton X-100, 0.5% Na-Desoxycholate) with added protease inhibitor cocktail (1:50) and protease/phosphatase inhibitor (1:50) and incubated for at least 10 min on ice. Lysates were mixed with SDS loading buffer (1:4) and were separated by SDS Page (10% acrylamide gel). Then transferred with a standard wet blot technique to a PVDF membrane. After 1 h blocking with 3% BSA in tween-tris buffered solution (TTBS) the proteins were detected with the antibody anti-CXCR7/RDC-1 (1:500) (Novus Biologicals, Centennial, USA) labeled with IRDyes and was incubated for 90min at room temperature in 3% BSA in TTBS. Equal loading was verified by actin (Abcam, Cambridge, UK) immunostaining. After drying the blots were detected by the Li-COR Odyssey System (LI-COR Biotechnology – GmbH, Bad Homburg, Germany) and analyzed by Li-COR Acquisition software (Version 5.2).

### Statistics and graphical presentation

Statistical analysis and the graphical data design were performed with GraphPad Prism (Graphpad Software, Inc., La Jolla, CA, USA, Version 10.1.1). For comparison of two sets normally distributed data a paired Student’s t-test was used. One-way ANOVA or Mixed-effects analysis were taken for more than two comparisons. Data are presented and given as Mean±SD. Multi-color flow cytometry data were analyzed and graphs designed by OMIQ flow cytometry software (Dotmatics, Boston, MA, USA).

## Results

### Hemin induces surface expression of ACKR3 on platelets

Previously, we found that expression of ACKR3 on the platelet plasma membrane is dynamically regulated upon activation [26, 27]. Platelet ACKR3 mediates antiapoptotic effects of platelets [28, 29] and loss of the receptor is associated with platelet hyperreactivity and apoptosis [24]. Recently, the hemoglobin degradation product hemin has been shown to induce platelet activation and non-apoptotic cell death by iron, known as ferroptosis [20, 21, 30]. To elucidate the effect of hemin on surface expression of ACKR3 on platelets, we stimulated isolated human platelets with hemin (6.25 or 25 µM) and surface expression of ACKR3 was determined using an anti-ACKR3 monoclonal antibody (anti-CXCR7 BV421). We found that hemin significantly enhances ACKR3 surface expression in a concentration-dependent manner (6.25 µM hemin: p□0.001; 25 µM hemin: p<0.0001) (**Figure 1A**). Total ACKR3 protein expression was not altered in response to hemin as demonstrated by western blot analyses (**Figure 1B**). Dynamic surface expression of ACKR3 was verified by image stream analysis (ImageStream^X^ Mark II Imaging Flow Cytometer, amnis, Seattle, WA, USA) (**Figure 1C**).

**Figure 1.**
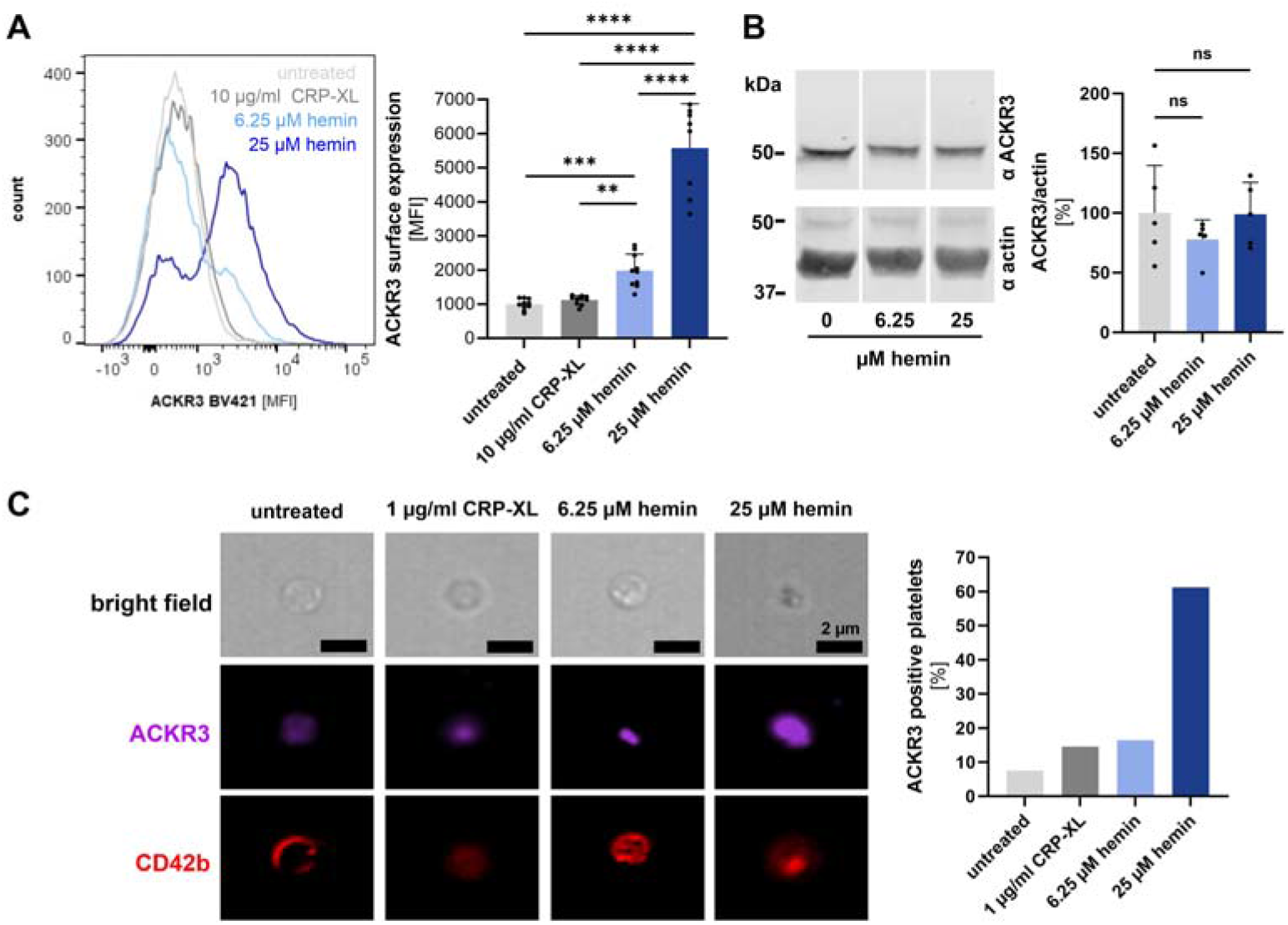
Hemin induces surface expression of ACKR3 on platelets. (A) Flow cytometry measurement of ACKR3 induced by 10 µg/ml CRP-XL, 6.25 and 25 µM hemin. Representative histogram (left) and statistical analysis (right); plotted: Mean ± SD; n = 10; statistics: Mixed-effects analysis, ^**^ p < 0.05, ^***^ p < 0.001, ^****^ p < 0.0001. (B) Representative blot and statistical analysis of total ACKR3 under 6.25 and 25 µM hemin stimulation compared to control conditions; plotted: Mean ± SD; n = 5; statistics: RM one-way ANOVA; ns not significant. (C) (left) Sample images of dynamic surface expression of ACKR3 captured by image stream analysis performed with isolated platelets at RT; scale bar = 2 µm. (right) Bar diagram of ACKR3 positive platelets measured with image stream.

### ACKR3 agonism attenuates hemin-induced platelet activation

Previously, we found that agonism of ACKR3 inhibits platelet activation and platelet-dependent thrombus formation [24, 25, 31]. To evaluate the effect of ACKR3 agonism on hemin-induced activation, we incubated isolated platelets with hemin (6.25 µM) and abundancy of activated fibrinogen receptor GPIIb-IIIa was determined using a conformation-dependent PAC-1 monoclonal antibody. We found that ACKR3 agonists (100 µM VUF11207, 100 µM BY-061) but not an inactive control chemical (100 µM C46) significantly attenuated hemin-dependent activation of GPIIb-IIIa (6.25 µM hemin vs. VUF11207 + 6.25 µM hemin: p<0.001; 6.25 µM hemin vs. BY-061 + 6.25 µM hemin: p<0.001) (**Figure 2A**). Further, platelet degranulation in response to hemin was significantly mitigated in the presence of specific ACKR3 agonists as shown by surface expression of P-selectin (CD62P, α-granula) (6.25 µM hemin *vs*. VUF11207 + 6.25 µM hemin: p<0.05; 6.25 mM hemin *vs*. BY-061 + 6.25 µM hemin: p<0.01) and CD63 (lysosomes) (6.25 µM hemin vs. VUF11207 + 6.25 µM hemin: p<0.05; 6.25 µM hemin *vs*. BY-061 + 6.25 µM hemin: p<0.05) (**Figure 2B and C**). Platelet degranulation is dependent on intracellular Ca^2+^ signaling [32, 33]. As described for platelet release, hemin induces a significant increase in intracellular calcium indicated by a rise Fluo4 fluorescence intensity (p<0.01) (**Figure 2D**). In presence of ACKR3 agonists hemin-dependent Ca^2+^ signaling was significantly attenuated (p<0.01) (**Figure 2D**).

**Figure 2.**
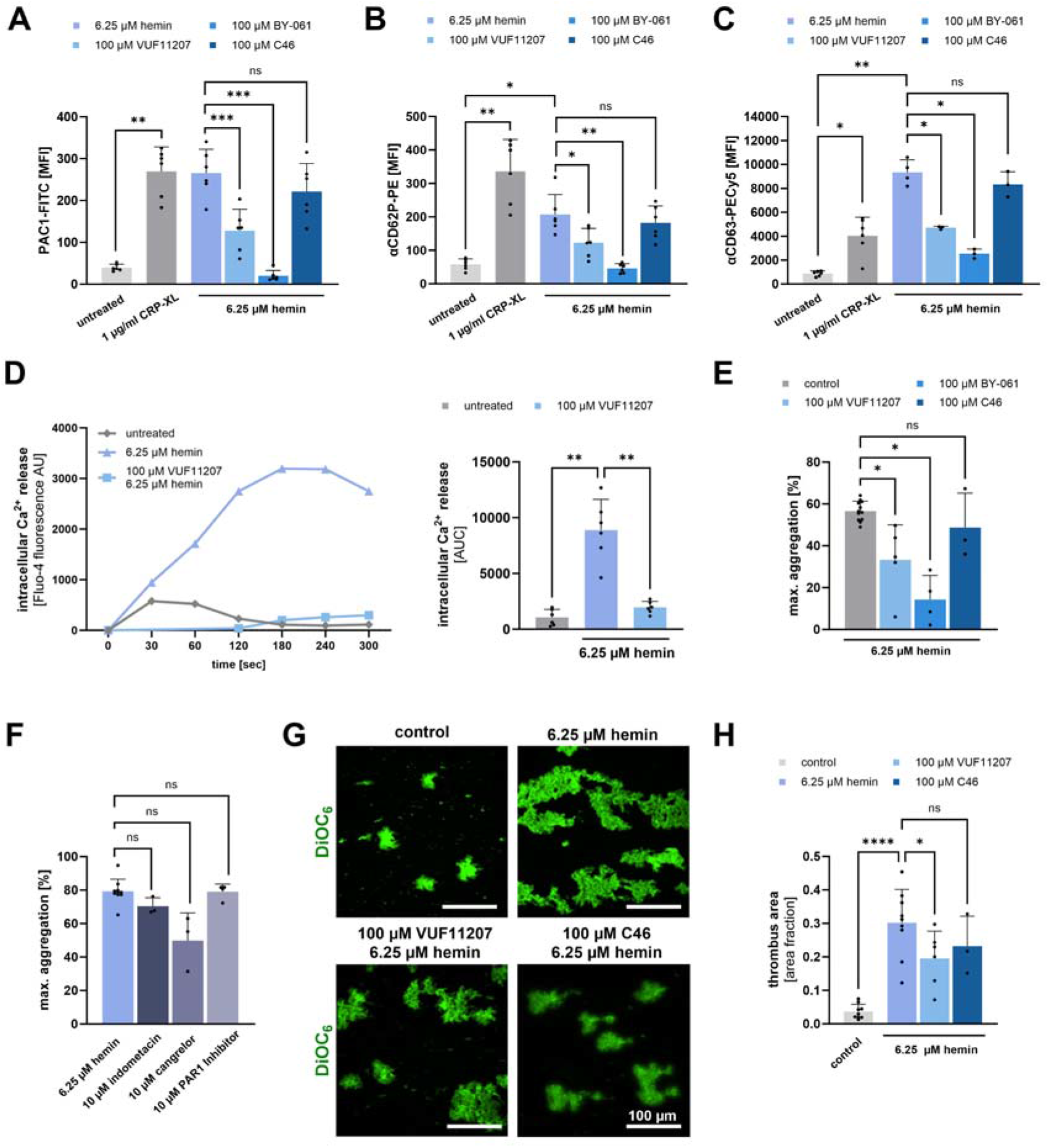
Effect of ACKR3 agonism on hemin-induced platelet activation, degranulation and thrombus formation. (A-C) Flow cytometry measurements of (A) PAC-1 (B) CD62P and (C) CD63 induced by 1 µg/ml CRP-XL and 6.25 µM hemin with pre-treatment of ACKR3 agonists (100 µM VUF11207, 100 µM BY-061, 100 µM C46) for 15 min at RT; plotted: Mean ± SD; n ≥ 3; statistics: RM one-way ANOVA (PAC1, CD62P), Mixed-effects analysis (CD63), ^*^p < 0.05; ^**^p < 0.01, ^***^p <0.001, ns not significant. (D) Representative intracellular Ca^2+^-release measurement for 5 min using Fluo-4 and statistical analysis of the area under the curve (AUC) of the Fluo-4 fluorescence; plotted: Mean ± SD; n = 6; statistics: Mixed-effects analysis, ^**^p<0.01 (E-F) Light transmission aggregometry measurements. (E) Hemin-induced maximal aggregation after 5 min at 37°C and pre-treatment with ACKR3 agonists (100 µM VUF11207, 100 µM BY-061, 100 µM C46) for 15 min at 37°C and a 6.25 mM hemin control, plotted: Mean ± SD; n ≥ 3; statistics: Mixed-effects analysis, ^*^p<0.05, ns not significant. (F) Hemin-induced maximal aggregation after 5 min at 37°C and pre-treatment with either COX-1 inhibitor (10 µM indometacin), P2Y_12_-inhibitor (10 µM cangrelor) or PAR1 inhibitor (10 µM ML161) for 15 min at 37°C and a 6.25 µM hemin control, plotted: Mean ± SD; n ≥ 3; statistics: Mixed-effects analysis, ns not significant. (G) Isolated human platelets were activated with 6.25 µM hemin and pre-treated with ACKR3 agonists (100 µM VUF11207, 100 µM C46) for 15 min at RT and perfused over a collagen-coated surface (100 µg/ml) at a shear rate of 500 s^-1^. Representative fluorescence microscopy images of thrombi stained with DiOC_6_; scale bar = 100 µm. (H) Diagram depicts thrombus area in the presence and absence of ACKR3 agonists with 6.25 µM hemin stimulation; plotted: Mean ± SD; n ≥ 3; statistics: Mixed-effects analysis, ^*^p <0.05, ^****^p <0.0001, ns not significant.

Next, we tested the effect of ACKR3 agonists on hemin-dependent platelet function. We found that agonism of ACKR3 inhibits hemin-dependent platelet aggregation in response to hemin (6.25 µM hemin *vs*. VUF11207 + 6.25 µM hemin: p<0.05; 6.25 µM hemin *vs*. BY-061 + 6.25 µM hemin: p<0.05) (**Figure 2E**). In contrast, and as shown previously [21], conventional antiplatelet compounds such as a COX-1 inhibitor (10 µM indomethacin), an ADP-receptor blocker (10 µM cangrelor), or a PAR1 inhibitor (10 µM ML161) did not inhibit platelet aggregation in response to hemin (**Figure 2F**). Similarly, in the presence of ACKR3 agonists hemin-dependent thrombus formation on immobilized collagen under flow was substantially reduced (p<0.05) (**Figure 2G**).

### ACKR3 agonism preserves integrity of platelets in the presence of hemin

Recently, we found that hemin leads to platelet death and plasma membrane fragmentation [21]. In the presence of ACKR3 agonists hemin stimulation did not result in substantial platelet metamorphosis (**Figure 3A**). Most strikingly, hemin-dependent fragmentation of the platelet plasma membrane was not observed (**Figure 3A**).

**Figure 3.**
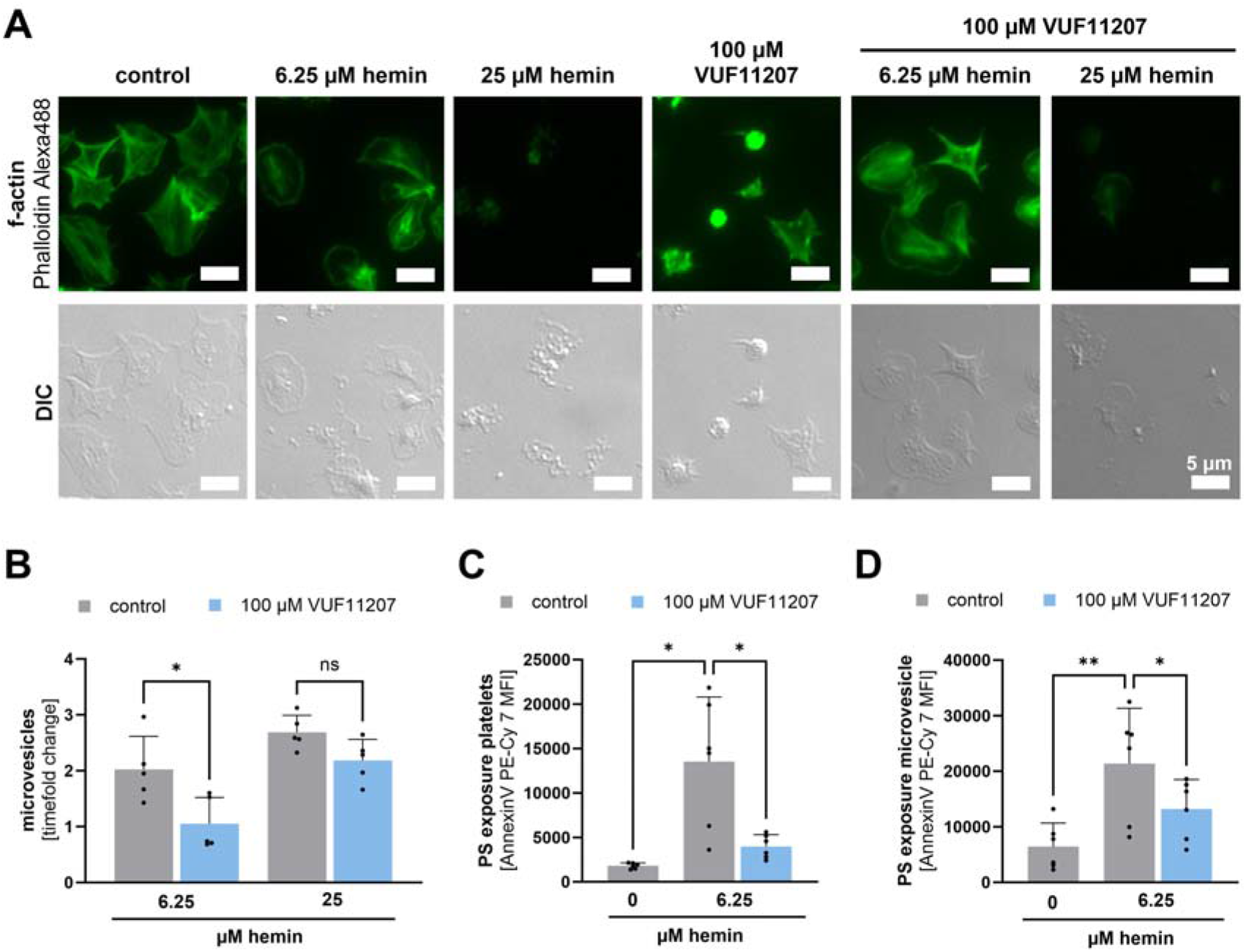
Effect of ACKR3 agonism on hemin-induced plasma fragmentation. (A) Sample images of spreaded platelets on 100 µg/ml fibrinogen coated coverslips with 6.25 and 25 µM hemin activation and pre-treatment with ACKR3 agonist (100 µM VUF11207) for 15 min at RT; bar scale = 5 µm. (B) Flow cytometry measurement of microvesicles (< 1 µm) with beads measured by BD FACSLyric™; plotted: Mean ± SD; n = 5; statistics: RM one-way ANOVA; ^*^p<0.05, ns not significant. (C/D) Flow cytometry measurement of phosphatidylserine (PS) exposure on platelets (C) and platelet-derived microvesicles (D) with an Annexin V conjugate (Annexin V PE-Cyanin7); plotted: Mean ± SD; n = 6; statistics: RM one-way ANOVA; ^*^p<0.05, ^**^p<0.01.

The formation of platelet fragments was also analyzed by flow cytometry showing that hemin induces concentration-dependent microvesicle formation (6.25 µM hemin: 2-fold change; 25 µM hemin: 2.7-fold change) and that ACKR3 agonist (100 µM VUF11207) reduced hemin-dependent formation of plasma membrane microvesicles (6.25 µM hemin vs. VUF11207 + 6.25 µM hemin: p<0.05) (**Figure 3B**). At high hemin concentrations (25 µM hemin) ACKR3 agonist does not decrease microvesicle formation (**Figure 3B**). The change of plasma membrane integrity was observed in response to hemin by an increase in phosphatidylserine exposure (PS) as indexed by enhanced Annexin V binding both on intact platelets (6.25 µM hemin vs. VUF11207 + 6.25 µM hemin: p<0.05) (**Figure 3C**) and on platelet-derived microvesicles (6.25 µM hemin *vs*. VUF11207 + 6.25 µM hemin: p<0.01) (**Figure 3D**). In the presence of ACKR3 agonists PS exposure both on platelets and platelet-derived microvesicles was reduced (platelets: 6.25 µM hemin *vs*. VUF11207 + 6.25 µM hemin: p<0.05) (**Figure 3C**) (microvesicles: 6.25 µM hemin *vs*. VUF11207 + 6.25 µM hemin: p<0.05) (**Figure 3D**).

### Platelet phenotype and formation of subpopulation is significantly altered in response to hemin and reversed by ACKR3 agonism

To get further insights into the effects of hemin on formation of platelet subpopulations, we established a 10-color flow cytometric assay including various monoclonal antibodies and dyes (anti-CD42b, PAC-1, anti-CD61, anti-CD41, anti-CD62P, anti-CD63, anti-CXCR4, anti-ACKR3, Annexin V and Zombie NIR) [34, 35]. Multi-color flow cytometry allows to define distinct platelet subpopulations associated with distinct function (e.g., aggregatory, procoagulant) [36-38]. We extended the multi-color FACS with the addition of antibodies against the chemokine receptors CXCR4 and ACKR3 (anti-CXCR4, anti-ACKR3). We found that hemin favors the formation of aggregatory (CD42b^pos^/ PAC-1^pos^/ AnnexinV^neg^) (p<0.0001), procoagulant (CD42b^pos^/ PAC-1^neg^/ AnnexinV^pos^) (p<0.0001) [21, 38] and a novel inflammatory (CD42b^pos^ / CXCR4^pos^/ ACKR3^pos^/ AnnexinV^pos^) platelet subpopulation (**Figure 4B and 4D**). The subpopulation pattern is significantly different in experiments when CRP-XL was used as platelet agonist (**Figure 4B**) which enhanced primarily or exclusively the aggregatory and procoagulant but not the inflammatory phenotype (**Figure 4A and 4D**). Most strikingly, in the presence of ACKR3 agonist (100 µM BY-061), formation of both aggregatory (6.25 µM hemin *vs*. BY-061 + 6.25 µM hemin: p<0.05) and procoagulant (6.25 µM hemin *vs*. BY-061 + 6.25 µM hemin: p<0.05) as well as inflammatory platelets (6.25 µM hemin *vs*. BY-061 + 6.25 µM hemin: p<0.05) was prevented (**Figure 4B and 4D**).

**Figure 4.**
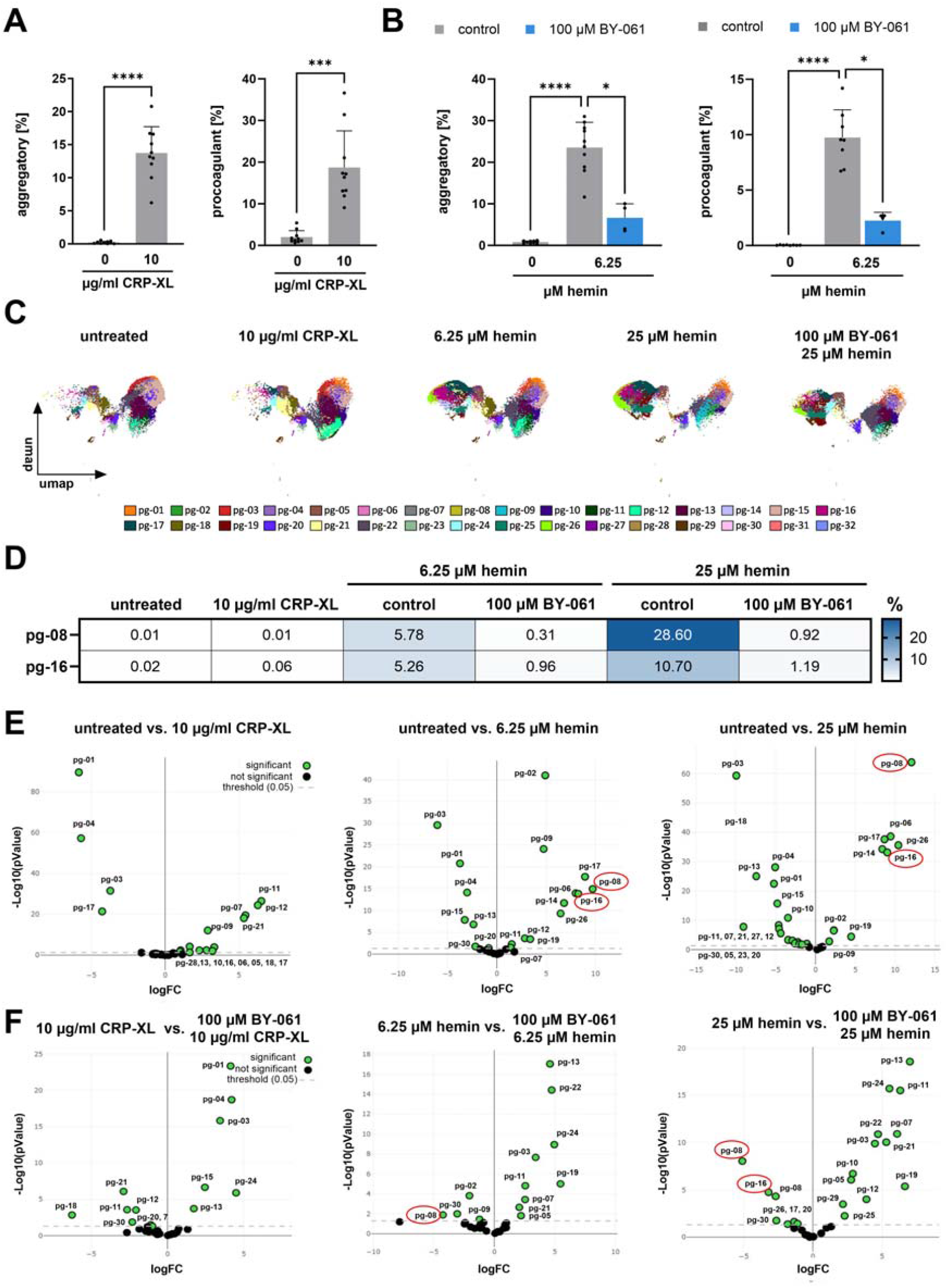
Platelet phenotype and formation of hemin-induced subpopulations. (A) Induction of aggregatory and procoagulant platelet subpopulation by 10 µg/ml CRP-XL measured by flow cytometry; plotted: Mean ± SD; n = 10; statistics: paired t-test, ^***^p<0.001, ^****^p<0.0001. (B) 6.25 µM hemin-induced aggregatory and procoagulant platelet subpopulations with 15 min pre-treatment of ACKR3 agonist (100 µM BY-061) measured with flow cytometry; plotted: Mean ± SD; n ≥ 4; statistics: Mixed-effects analysis, ^*^p<0.05, ^****^p<0.0001. (C) Platelet subpopulations determined by PhenoGraph algorithm for unsupervised clustering (pg-01 to pg-32) of human platelets; n = 10. Plots represent and overlay of all platelets per treatment (untreated, 10 µg/ml CRP-XL, 6.25 µM hemin, 25 µM hemin, 100 µM BY-061 + 25 µM hemin). (D) Inflammatory subpopulations. Abundancy of platelets in each inflammatory cluster induced by 10 µg/ml CRP-XL, 6.25 and 25 µM hemin with pre-treatment of ACKR3 agonist (100 µM BY). (E) Volcano plot presentation of platelet subpopulations in untreated versus treatment samples (10 µg/ml CRP-XL, 6.25 µM hemin, 25 µM hemin); p<0.05. (F) Volcano plot diagram subpopulations generated by treatment versus treatment with pre-incubation of ACKR3 agonist (100 µM BY-061); threshold: p<0.05.

To further define the effect of hemin on the formation of platelet subpopulations, we performed unsupervised data analysis by applying uniform manifold approximation and projection (UMAP) dimension reduction to group phenotypically similar events followed by unsupervised clustering analysis using PhenoGraph (**Figure 4C**). PhenoGraph analysis resolved 32 clusters (pg-01 to pg-32) (**Figure 4C**), of which only two (pg-08 and pg-16) occurred under hemin-treatment and showed significant differences between untreated and hemin-treated platelets depending on the hemin concentration used (**Figure 4D**). At low hemin concentrations (6.25 µM) 5.78% of pg-08 and 5.26% of pg-16 appear. At higher hemin concentrations (25 µM) both pg-08 and pg-16 increase (pg-08: 28.6%; pg-16: 10.7%) (**Figure 4D**). Most strikingly, ACKR3 agonism (100 µM BY-061) attenuated primarily the two clusters (pg-08 and pg-16) of hemin-induced formation of platelet subpopulations with high expression levels of Annexin V (procoagulant) and CXCR4/ACKR3 (inflammatory) (BY-061 + 6.25 µM hemin: pg-08 (0.31%), pg-16 (0.96%); BY-061 + 25 µM hemin: pg-08 (0.92%), pg-16 (1.19%)) (**Figure 4D**). The volcano plot confirms the significant (p<0.05) increase in inflammatory subpopulations (CD42b^pos^/ CXCR4^pos^/ ACKR3^pos^/ AnnexinV^pos^) pg-08 and pg-16 and the reduction of these populations by ACKR3 agonisms (**Figure 4E and 4F**).

## Discussion

The major findings of the present study are: **(i)** Hemin induces surface expression of the chemokine receptor ACKR3 on platelets. **(ii)** ACKR3 agonism inhibits hemin-induced platelet activation, degranulation, thrombus formation and plasma membrane fragmentation. **(iii)** Hemin-dependent changes of the platelet phenotype and formation of aggregatory and procoagulant subpopulations are significantly altered through ACKR3 agonism. Our data indicate that iron-overload due to Fe^3+^ and Cl^-^ -containing hemin results in substantial platelet-dependent thrombus formation *ex vivo* and formation of procoagulant platelet subtypes. Agonistic enhancement of the chemokine receptor ACKR3 may be a novel and promising strategy to limit platelet activation and thrombus formation at site of intravascular hemorrhages that are associated with enhanced vulnerability of atherosclerotic plaques. The occurrence of intraplaque hemorrhages is well described in atherosclerotic plaques [8-12, 14]. Besides lipid-rich vascular areas the presence of intraplaque hemorrhages (IPH) promotes destabilization and vulnerability of an atherosclerotic plaque resulting in uncontrolled intraluminal thrombus formation and acute vessel occlusion leading to myocardial infarction or ischemic stroke [11, 12, 14]. IPH are caused by ruptured neovessels [9, 41] and are characterized by release extracellular hemoglobin and Fe^3+^ and Cl^-^-containing hemin [12, 42]. Only recently, hemin has been identified as a strong platelet activating factor leading to platelet-dependent thrombus formation and substantial plasma membrane receptor surface expression [20-22]. Interestingly, conventional anti-platelet drugs such as COX1 inhibitors (indomethacin) or ADP-receptor blocker have only modest if at all inhibitory effects on hemin-induced platelet activation [21]. This indicates that the activation signals in response to hemin are different compared to classical COX-1- or P2Y_12_-dependent activation mechanisms. Recently, we described that modulation of cGMP levels or inhibition of the subtilisin-like proprotein-processing enzyme furin attenuates hemin-induced platelet activation [21, 22].

Hemin-induced platelet activation substantially induce non-apoptotic iron-mediated cell death, called ferroptosis which has been well documented in nuclear cells [43]. In platelets, hemin induces significant alterations of phosphatidylserine (PS) exposure on the plasma membrane, elevation of reactive oxygen species (ROS) and a loss of mitochondrial membrane potential [26, 44]. This indicates that hemin induces platelet-cell death at site of bleeding or IPH.

Previously, we found that the noncanonical chemokine receptor ACKR3 (formally CXCR7) is an inhibitory receptor for platelet activation [23, 24]. Selective ACKR3 agonists are strong inhibitors of platelet activation and thrombus formation [23, 31, 45]. The fact that classical platelet antagonists do not substantially attenuate platelet activation or PS exposure in response to hemin encouraged us to study the effect of ACKR3 agonists in this context. We found that hemin enhances surface expression of ACKR3 which has been described as survival receptor in platelets. Agonistic enhancement of ACKR3 in response to agonists substantially mitigated platelet activation and platelet-dependent *ex vivo* thrombus formation under flow, an effect that was not detected in the presence of COX-1, P2Y_12_ or PAR1 antagonists [21].

Interestingly, ACKR3 agonists significantly suppressed the formation of procoagulant platelet subtypes and membrane microvesicles, which play a major role in to triggering coagulation in the surrounding of platelet accumulation. Thus, it is tempting to speculate that at a site of tissue bleeding or IPH an enhanced hemin concentration might result in increased platelet activation and coagulation activity which in turn might contribute to limit bleeding. Further, uncontrolled hemin-dependent platelet activation and procoagulant activity may result in vulnerability of an atherosclerotic plaque and enhancing the risk for intravasal thrombus formation and occurrence of myocardial infarction or ischemic stroke. At present, we do not have evidence that our hypothesis holds true for the *in vivo* situation. An adequate *in vivo* model is not easily established since systemic administration of hemin is rapidly inhibited by the presence of abundant plasmatic protein including hemopexin, haptoglobin or albumin [46-48]. This limits our translational significance of our hypothesis. However, since conventional preventive antithrombotic drugs for patients at risk for atherothrombotic events may not be an optimal strategy to limit vulnerability of plaques, the hypothesis of ACKR3 agonism may be a promising concept that needs to be developed.

## Abbreviations

ACD: acid-citrate-dextrose
ACKR3: atypical chemokine receptor 3 (formally CXCR7)
ADP: adenosine diphosphate
ANOVA: analysis of variance
BSA: bovine serum albumin
CD41: classification determinant 41/integrin α-2b
CD42b: classification determinant 42b/glycoprotein-Ib
CD61: classification determinant 61/integrin β3
CD62P: classification determinant 62P/P-Selectin
CD184: classification determinant 184/C-X-C chemokine receptor type 4
CLEC-2: C-type lectin-like type-2
cGMP: cyclic guanosine monophosphate
CXCR4: C-X-C chemokine receptor type 4
COX-1: cyclooxygenase 1
CRP-XL: collagen related peptide
DiOC_6_: 3,3’-Dihexyloxacarbocyanine Iodide
DIC: interference contrast microscopy
FSC: forward scatter
GPIIb-IIIa: glycoprotein IIb-IIIa
GPVI: glycoprotein VI
Hb: hemoglobin
IPH: intraplaque hemorrhage
P2Y_12_: chemoreceptor for adenosine diphosphate
PAR1: protease activated receptor 1
PBS: Dulbecco’s phosphate buffered solution
PS: phosphatidyl serine
PVDF: polyvinylidene difluoride
RBC: red blood cell
RIPA: radioimmunoprecipitation assay
RM: repeated measures
ROS: reactive oxygen species
RT: room temperature
SD: standard deviation
SDS: sodium dodecyl sulfate
SDS-PAGE: sodium dodecyl sulfate-polyacrylamide gel electrophoresis
SSC: side scatter
TTBS: tween-tris buffered solution
TRIS: tris(hydromethyl)aminomethane
UMAP: uniform manifold approximation and projection

## Author Contributions

Experiment design and planning: M.G; Z.L..; V.D.-B.; D.S.; A.-K.R; experiment performance: Z.L.; V.D.-B.; D.S.;R.H.; A.B; data analysis and plotting: Z.L.; V.D.-B.; D.S.; R.H. T.H.; interpretation and discussion of results: Z.L.; V.D.-B.; D.S.; R.H.; M.S., T.H., T.C.; A.-K.R.; M.G. figure design: Z.L.; M.G.; manuscript writting: M.G.; Z.L.; T.C.

## Acknowledgements

This project was supported by the German Research Foundation (DFG) – Project number 335549539 – GRK 2381. David Schaale was supported by a research grant of the German Cardiac Society (DGK).

## Conflicts of Interest

The authors declare no competing financial interests.

## Notes

### Competing Interest Statement

The authors have declared no competing interest.

### Summary of Updates

We made a mistake in the surname of one of the coauthors. "Anne-Katrin Rohfling" was corrected to "Anne-Katrin Rohlfing".

